# The evolution of genomic, transcriptomic, and single-cell protein markers of metastatic upper tract urothelial carcinoma

**DOI:** 10.1101/2021.11.16.468622

**Authors:** Kentaro Ohara, André Figueiredo Rendeiro, Bhavneet Bhinder, Kenneth Wha Eng, Hiranmayi Ravichandran, David Pisapia, Aram Vosoughi, Evan Fernandez, Kyrillus S. Shohdy, Jyothi Manohar, Shaham Beg, David Wilkes, Brian D. Robinson, Francesca Khani, Rohan Bareja, Scott T. Tagawa, Andrea Sboner, Olivier Elemento, Bishoy M. Faltas, Juan Miguel Mosquera

**Affiliations:** Department of Pathology and Laboratory Medicine, Weill Cornell Medicine, New York, NY 10065, USA; Englander Institute for Precision Medicine, Weill Cornell Medicine, New York, NY 10021, USA; Department of Physiology and Biophysics, Weill Cornell Medicine, 1300 York Avenue New York, NY 10065, USA; Institute for Computational Biomedicine, Weill Cornell Medicine, 1305 York Avenue, New York, NY, 10021, USA; Sandra and Edward Meyer Cancer Center at Weill Cornell Medicine, New York, NY 10065, USA; Department of Medicine, Division of Hematology and Medical Oncology, Weill Cornell Medicine, New York, NY 10065, USA; Department of Cell and Developmental Biology, Weill Cornell Medicine, New York, NY 10065, USA

## Abstract

The molecular characteristics of metastatic upper tract urothelial carcinoma (UTUC) are unknown. The genomic and transcriptomic differences between primary and metastatic UTUC is not well described either. We combined whole-exome sequencing, RNA-sequencing, and Imaging Mass Cytometry^™^ (IMC^™^) of 44 tumor samples from 28 patients with high-grade primary and metastatic UTUC. IMC enables spatially resolved single-cell analyses to examine the evolution of cancer cell, immune cell, and stromal cell markers using mass cytometry with lanthanide metal-conjugated antibodies. We discovered that actionable genomic alterations are frequently discordant between primary and metastatic UTUC tumors in the same patient. In contrast, molecular subtype membership and immune depletion signature were stable across primary and matched metastatic UTUC. Molecular and immune subtypes were consistent between bulk RNA-sequencing and mass cytometry of protein markers from 340,798 single-cells. Molecular subtyping at the single cell level was highly conserved between primary and metastatic UTUC tumors within the same patient.

## Introduction

Upper tract urothelial carcinoma (UTUC) is defined as urothelial carcinoma (UC) arising from the ureter or the renal pelvis. UTUC is a rare tumor compared to UC of the bladder (UCB), accounting for 5-10% of UC, with aggressive clinical presentation. While 5-year survival for non-muscle invasive UTUC is more than 90%, it drops to < 40% in patients with regional nodal metastases and < 10% in patients with distant metastases^1,2^. Even though systemic chemotherapies are given after relapsing, most patients die from the disease within 3 years^3^. A better understanding of the underlying molecular basis of metastatic UTUC is needed to develop targeted therapeutic strategies and improve clinical outcomes for patients with metastatic UTUC.

Recent studies by next-generation sequencing (NGS) have detailed the genomic landscape of primary UTUC^4-6^. Regarding molecular and immune phenotyping of UTUC, our group recently demonstrated that primary UTUC has a luminal-papillary T-cell depleted phenotype and a lower total mutational burden than UCB^7^. However, previous studies on UTUC including ours mainly focused on primary tumors and limited molecular data currently exists for metastatic UTUC^6^. Furthermore, the extent of genomic and phenotypic heterogeneity in each individual patient with metastatic UTUC is still unknown.

In this study, we performed whole-exome sequencing (WES), RNA-sequencing (RNA-seq), and multiplexed imaging cytometry which leverages antibodies conjugated to rare metals and detection by mass spectrometry to provide spatial protein expression patterns at a single cell resolution^8,9^ from patients with primary and metastatic UTUC enrolled in our precision medicine study to characterize the genomic, transcriptomic and immunophenotypic features of metastatic UTUC, and understand the degree of heterogeneity between primary and metastatic UTUC.

## Results

### Genomic landscape of metastatic UTUC

To explore the genomic landscape of metastatic UTUC, we performed WES of 44 prospectively collected UTUC tumors from a new cohort of 28 patients, including 7 matched sets of primary and metastatic UTUC and germline samples and 1 rapid autopsy (**Figure 1a, 1b** and **Supplementary Table. 1**). Overall, metastatic lesions showed an average of 196 non-synonymous SNVs/Indels (range 5 - 713) and 149 CNAs (amplification and deletion, range 4 – 831). All the analyzed UTUC had TMB scores below the UC-specific threshold we previously identified^10^ to designate TMB-high tumors. There were no significant differences in the MSIsensor scores^11^ and TMB between primary and metastatic UTUC (Mann-Whitney U-test *P* value = 0.65 and 0.22, respectively) (**Supplementary Figure 1**). The most frequently altered genes in our UTUC cohort were *TP53* (45.2%), *KMT2D* (35.7%), *ARID1A* (33.3%) and *CDKN2A* (33.3%) (**Figure 2a**). The frequently altered genes in our cohort were in agreement with previously reported primary UTUC genomic studies^4,5,7,12^, except for *FGFR3* (4/42, 9.5%). Of the detected CNAs, the frequency of *RAF1* amplification was significantly higher in metastatic UTUC (33.3%) compared to primary UTUC (4.2%) (Fisher exact test *P* value = 0.03) (**Figure 2b**).

**Figure 1.**
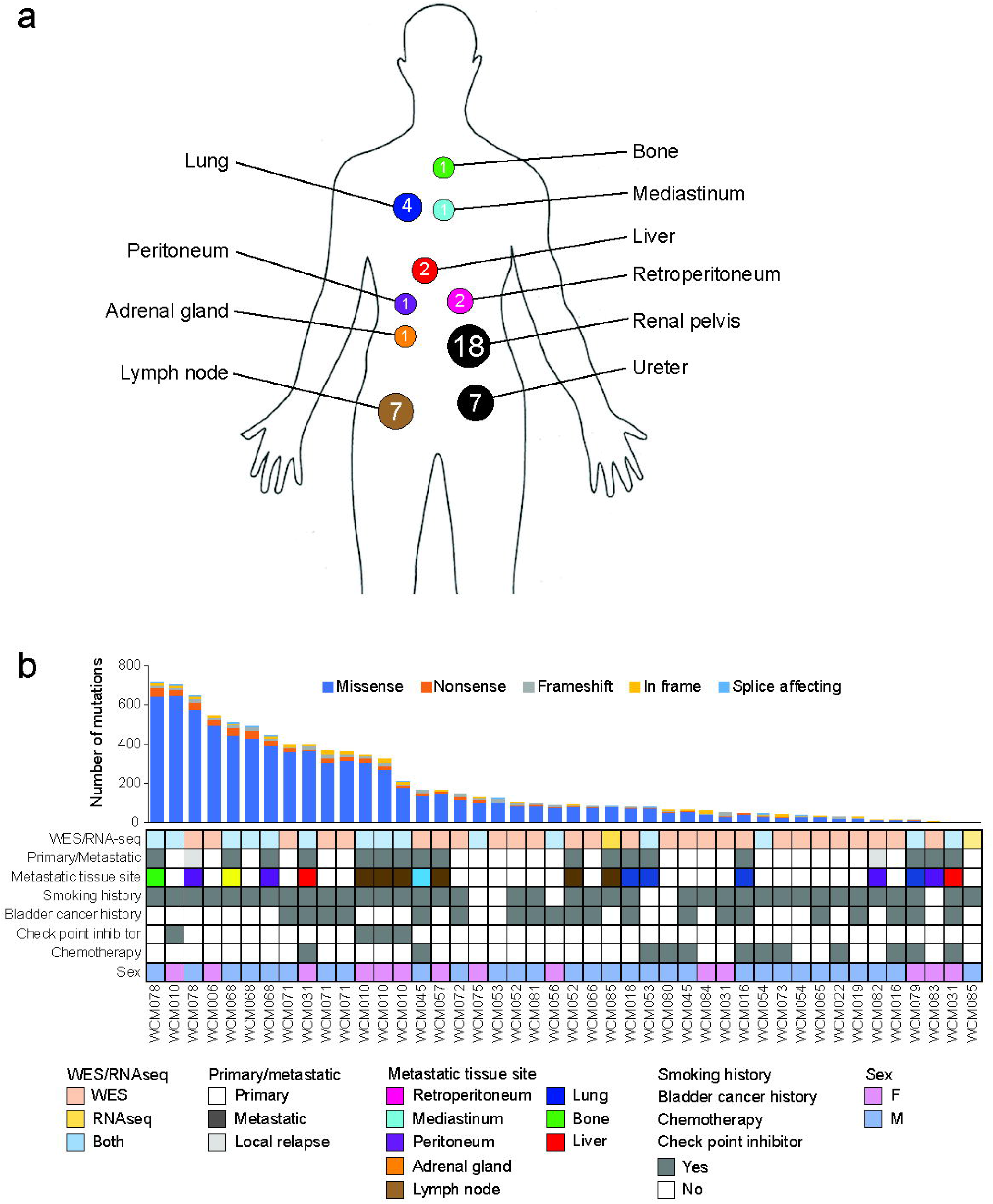
Clinical characteristics of the study cohort. **(a)** Schematic illustrating the anatomical sites of primary and metastatic upper tract urothelial carcinoma (UTUC) samples. Numbers correspond to the number of tumors at each site. **(b)** Clinical characteristics of the study cohort, the anatomical sites of primary and metastatic tumor samples and sequencing methods performed for each sample. Barplots represent numbers of non-synonymous mutations for each sample.

**Figure 2.**
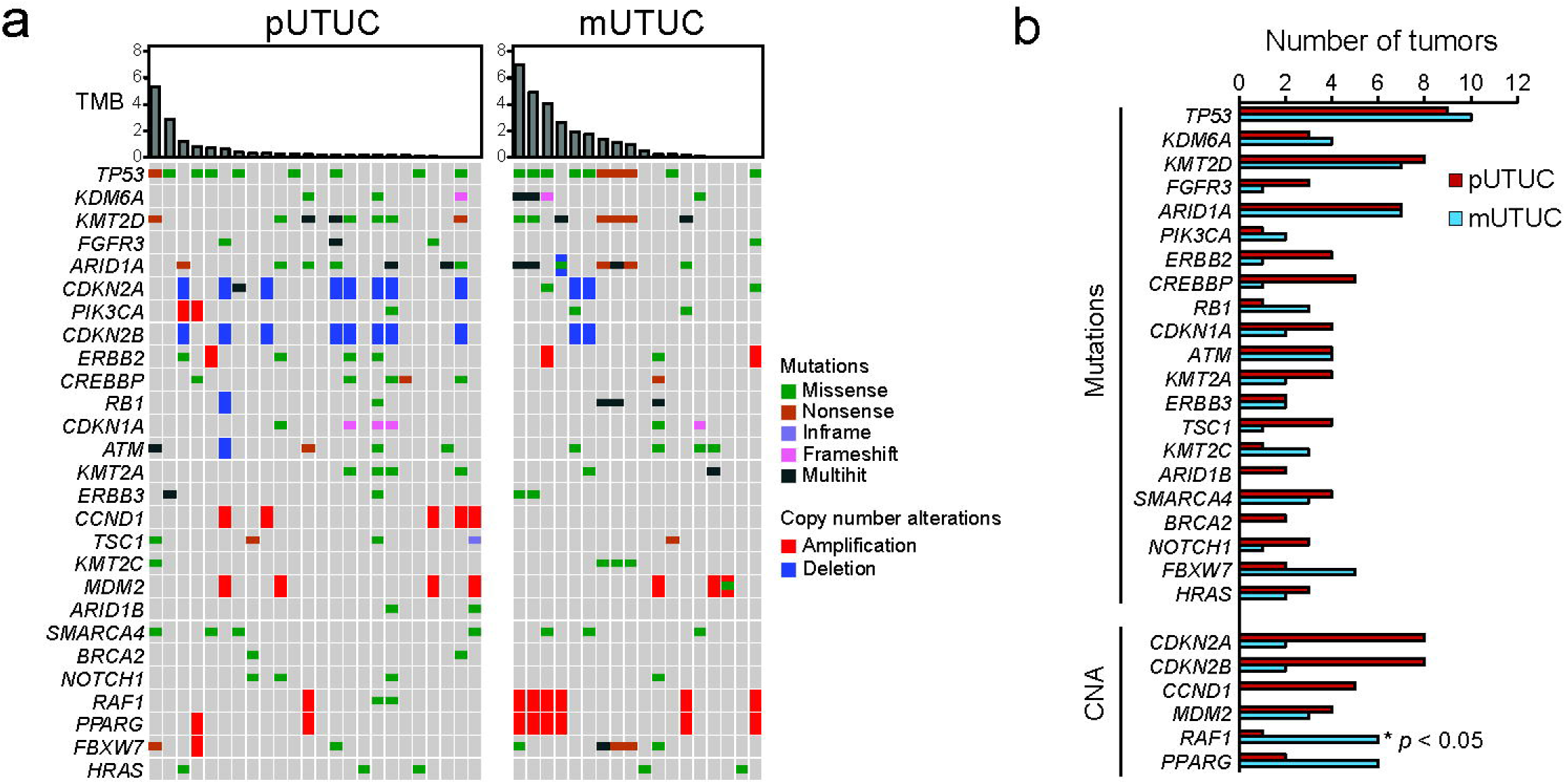
Genomic landscape of primary and metastatic UTUC. **(a)** Comparison of genomic alterations between primary and metastatic UTUC. **(b)** Comparison of genomic alteration frequencies between primary (red bars) and metastatic UTUC (blue bars) for genes frequently altered in our cohort.

### Genomic heterogeneity between primary and matched metastatic UTUC

To compare the clonal structure of primary and metastatic UTUC, we investigated the numbers of private and shared mutations between the primary and metastatic UTUC within each patient by calculating the percentage of non-synonymous mutations that were private to the primary tumor, the metastatic tumor or shared within individual patients. Of the 7 analyzed patients with paired primary and metastatic UTUC, 4 patients received chemotherapy treatment before sampling of the primary or metastatic tumor tissue, and all the tumors from two patients, WCM052 and WCM068, were chemotherapy-naïve (**Supplementary Figure 2**). On average, only 17.6% (range 7.3–36.4%) of mutations were shared by primary and matched metastatic samples (**Figure 3a**). The percentages of shared mutations were higher in the chemotherapy naïve patients than patients who received prior chemotherapy treatment (32.6% vs. 11.5%), suggesting that chemotherapy potentially plays a role in increasing genomic heterogeneity by inducing mutations^13-15^.

**Figure 3.**
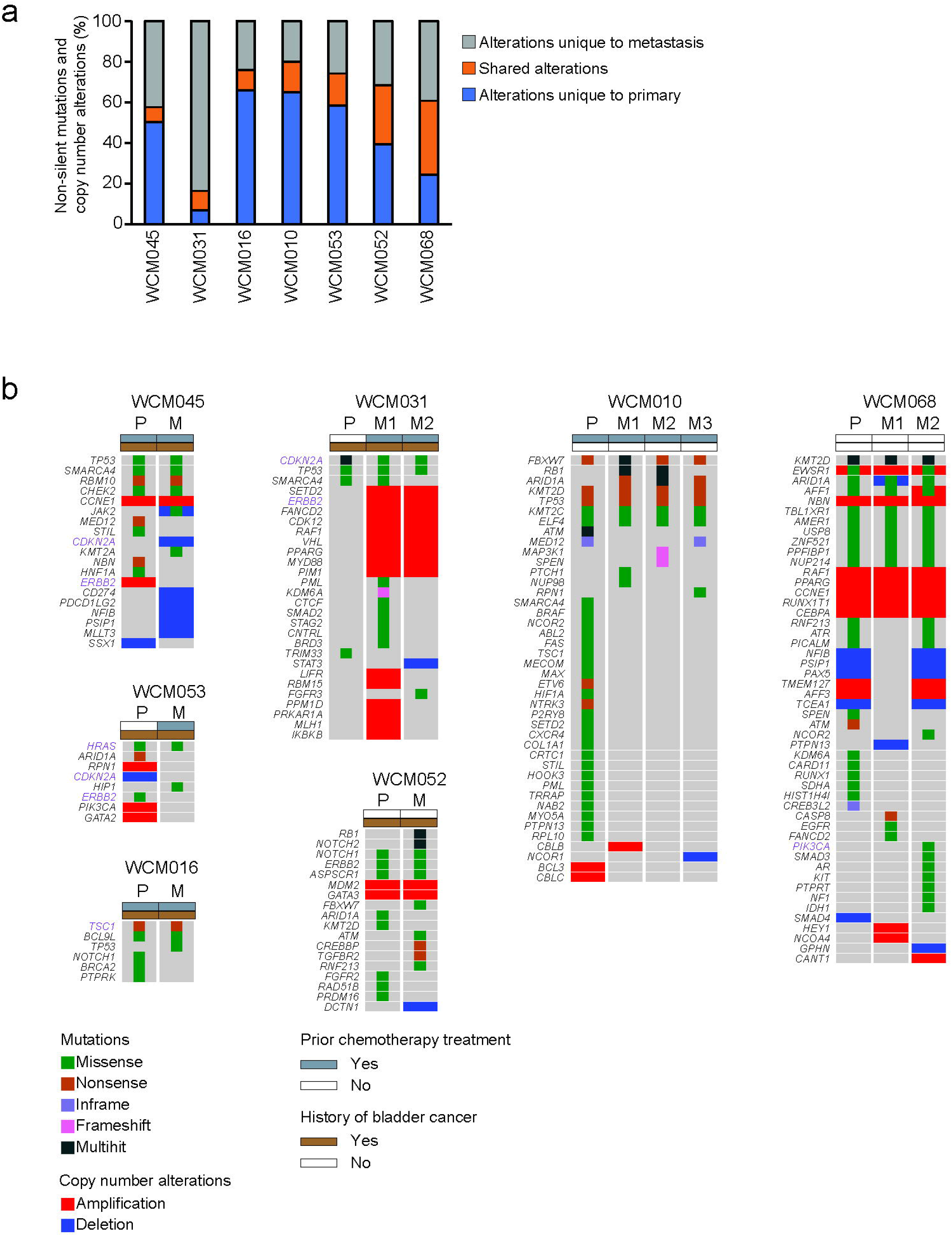
Genomic heterogeneity between paired primary and metastatic UTUC. **(a)** Percentages of somatic mutations unique to primary (blue) and metastasis (gray) or shared between the matched tumors (orange). **(b)** Oncoprints representing somatic mutations and copy number alterations of cancer genes in primary (P) and matched metastatic (M) UTUC. Oncoprints are grouped per case. Chemotherapy treatment prior to tissue collection and a history of bladder cancer are shown on the top of each oncoprint. Alterations shown in purple indicate actionable alterations.

To determine the biological impact of genomic heterogeneity, we next sought to define the differences in mutations and CNAs affecting cancer associated genes between primary and matched metastatic UTUC within each patient. A median of 13.5 cancer gene alterations per sample was identified (range 2 - 51), including a median of 2 (range 1–9) alterations predicted to be likely oncogenic by the OncoKB database^16^. When we compared genomic alterations between paired primary and metastatic tumors, intra-patient heterogeneity in cancer gene alterations was observed (**Figure 3b**). The primary and matched metastatic tumors shared at least one pathogenic cancer gene alteration in all cases (**Supplementary Figure 3**). However, compared to primary UTUC, we identified additional pathogenic mutations or CNAs in all but one of the matched metastatic UTUC tumors (**Figure 3b, Supplementary Figure 3**).

### Stability of molecular subtype and immune contexture assignments between primary and metastatic UTUC tumors

The next step was to see if there was any transcriptome difference between the primary and matched metastatic UTUC. To that goal, we performed whole-transcriptome sequencing (RNA-seq) on 17 UTUC tumors (six primary and eleven metastatic), including 3 patients with primary and matched metastatic UTUC and 1 patient with two metastatic tumors (**Figure 4a**). Using the recently published single sample consensus molecular classifier^17^, we found that 83.3% of primary tumor samples were luminal-papillary, and the rest were basal/squamous (**Figure 4a**). Of the metastatic tumors, 45.5%, 27.3%, 9.1% and 18.2% were luminal-papillary, luminal-unstable, stroma-rich and basal/squamous, respectively (**Figure 4a**). Clustering analysis using the BASE47 gene classifier^18^ showed a similar frequency of basal subtype between primary and metastatic tumors (33.3% vs. 27.3%, *P value* > 0.05). When we evaluated the molecular subtypes between primary and the matched metastatic UTUC tumors, we found that they were relatively stable in comparison to genomic changes, but discordance in molecular subtype was observed in one out of three cases (**Figure 4d**). In WCM010, one of three lymph node metastases was classified as luminal-unstable, while the primary and remaining lymph node metastases were classified as luminal-papillary. In case WCM031, only metastatic tumors were interrogated by RNA-seq due to the unavailability of frozen tissue from the primary tumor. Interestingly. molecular subtyping revealed discordance of molecular subtypes between the asynchronous liver metastases (**Figure 4b**). The initial metastasis (M1) and the second metastasis (M2) demonstrated stroma-rich and luminal unstable, respectively.

**Figure 4.**
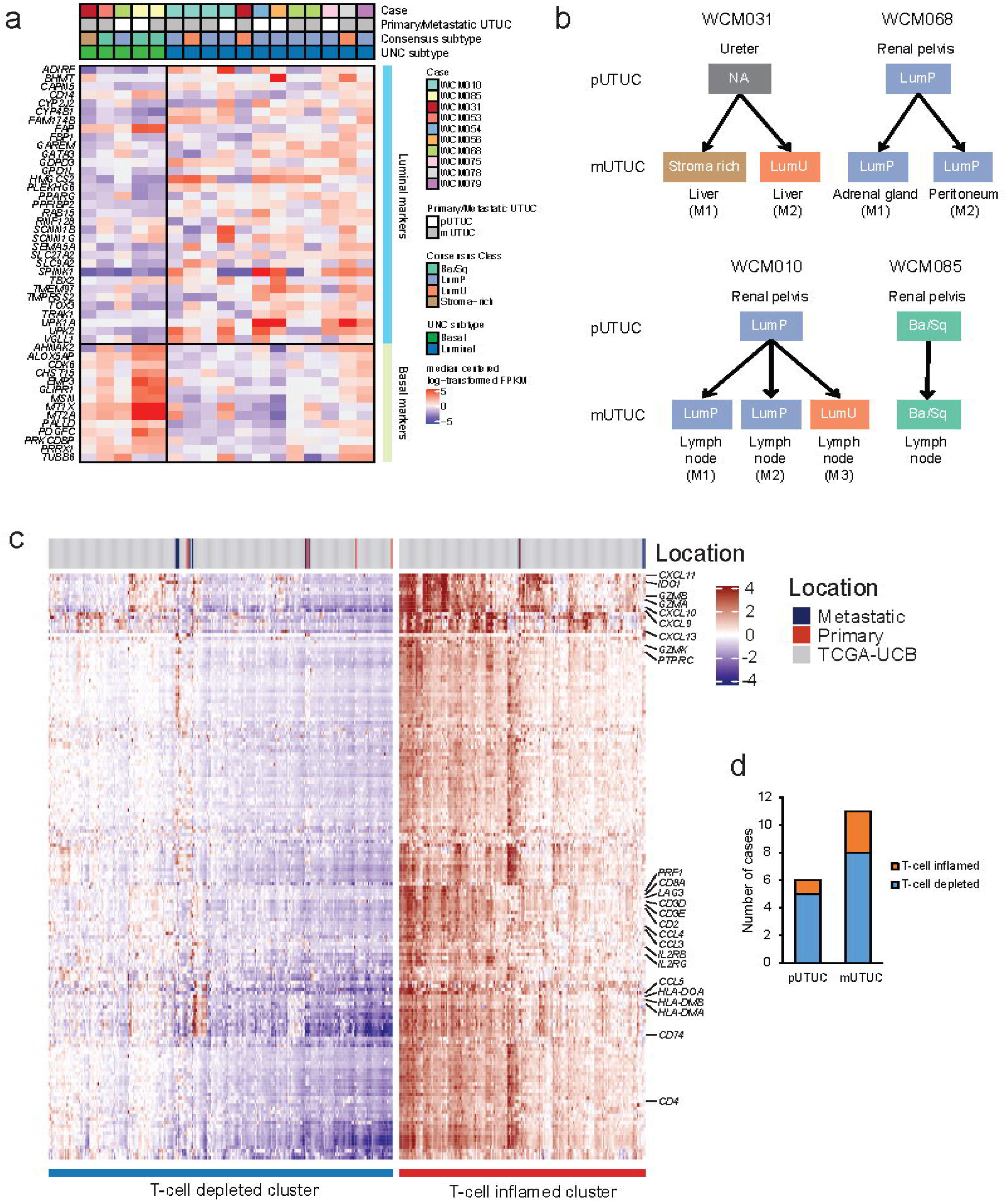
Comparison of molecular subtype and T-cell inflammation classification in primary and metastatic UTUC. **(a)** Gene expression heatmap of UTUC aligned by cases. The expression value is log-transformed and median centered for selected genes, for labeled gene sets. Assigned molecular classes are represented on top. The subtypes are labeled as luminal papillary (LumP), luminal non-specified (LumNS), luminal unstable (LumU) and basal/squamous (Ba/Sq). **(b)** Comparison of molecular subtypes between paired primary/metastatic and metastatic/metastatic UTUC. **(c)** Supervised consensus clustering of our UTUC and TCGA UCB tumors according to a 170-immune gene signature classifies tumors into T-cell depleted (with lower expression of classifier genes), and T-cell inflamed (with higher expression of classifier genes) clusters. **(d)** Breakdown of assigned classification for primary and metastatic UTUC.

Next, we applied a classifier of 170 immune-related genes which we previously developed^7^ (**Supplementary Table 2**) to the RNA-seq data obtained from 17 samples (6 primary and 11 metastatic UTUC) and the TCGA-BLCA cohort. The majority of UTUC tumors (76.5%, 13/17) clustered into the T-cell depleted subgroup (**Figure 4c-d**) consistent with our previous finding in primary UTUC^7^. Only 16.7% of primary UTUC (1/6) and 27.3% of metastatic UTUC (3/11) were T-cell inflamed. The immune phenotypes were concordant within individual patients.

### Single cell spatial profiling of UTUC’s tumor-immune contexture reveals intra-tumoral plasticity

In order to validate our findings showing differential immune states within the tumors and further investigate the tumor microenvironment, we employed Imaging Mass Cytometry^™^ (IMC^™^) on samples of UTUC – a method for multiplexed tissue imaging using antibodies conjugated to rare metals and detection by mass spectrometry^19^. In total, from 14 samples of 6 patients, we generated 58 images containing the spatial distribution of 27 distinct molecular biomarkers, which included markers of tumor cells (pan-Keratin, Keratin 5, GATA3, E-cadherin), immune cells (CD3, CD16, CD68), among others such as functional or stromal markers present in various cell types (**Supplementary Table 3**). Upon inspection, IMC^™^ images generally revealed structured boundaries between tumor and stromal compartments (**Figure 5a-b**). For a quantitative description of tumor and tumor-associated phenotypes, we segmented a total of 340,798 single-cells across all images and using the marker intensity as a quantitative measure of epitope abundance, built a joint space reflecting the phenotypic similarity between cells using the Uniform Manifold Approximation and Projection method (UMAP) (**Figure 5c**). In this space, the tumor cells were readily distinguishable from the remaining cells through the expression of the basal marker Keratin 5, or expression of either the luminal marker GATA3 or other keratin proteins (**Figure 5d**). The remaining cells formed the immune and stromal compartments of the tumor, of which the most numerous were cancer-associated fibroblasts, macrophages, CD4 and CD8+ T-cells, and endothelial cells (**Figure 5d-e**). We found that the samples classified as immune-inflamed by the RNA-seq based classifier had a significantly higher number T-cells with CD8 protein expression, effectively validating 170 gene RNA-seq based classifier for immune characterization of UTUC (**Figure 5f**). In the tumor compartment, we identified a wide range of tumor phenotypes expressing various combinations of these markers, including in the expression of tumor-specific proteins (**Figure 5e**). Upon closer inspection, we observed that the degree of plasticity as measured by the ratio of the expression of the basal marker KRT5 to the luminal marker GATA3 (KRT5/GATA3) in each single cell was largely conserved across primary and metastatic UTUC within the same patient (**Figure 5g**). However, for one patient (WCM031), the primary and metastatic tumor cells had distinct phenotypes, with the primary tumor having a basal phenotype with low CD8^+^ T-cell infiltration, while the metastatic sample has a luminal phenotype and approximately double the number of CD8^+^ T-cells (78.6 cells in primary vs 134.5 cells in metastasis, **Supplementary figure 4**). These findings suggest that cancer-immune contexture phenotypes are heterogeneous between patients but often conserved between primary and metastatic tumors within individual patients with UTUC.

**Figure 5.**
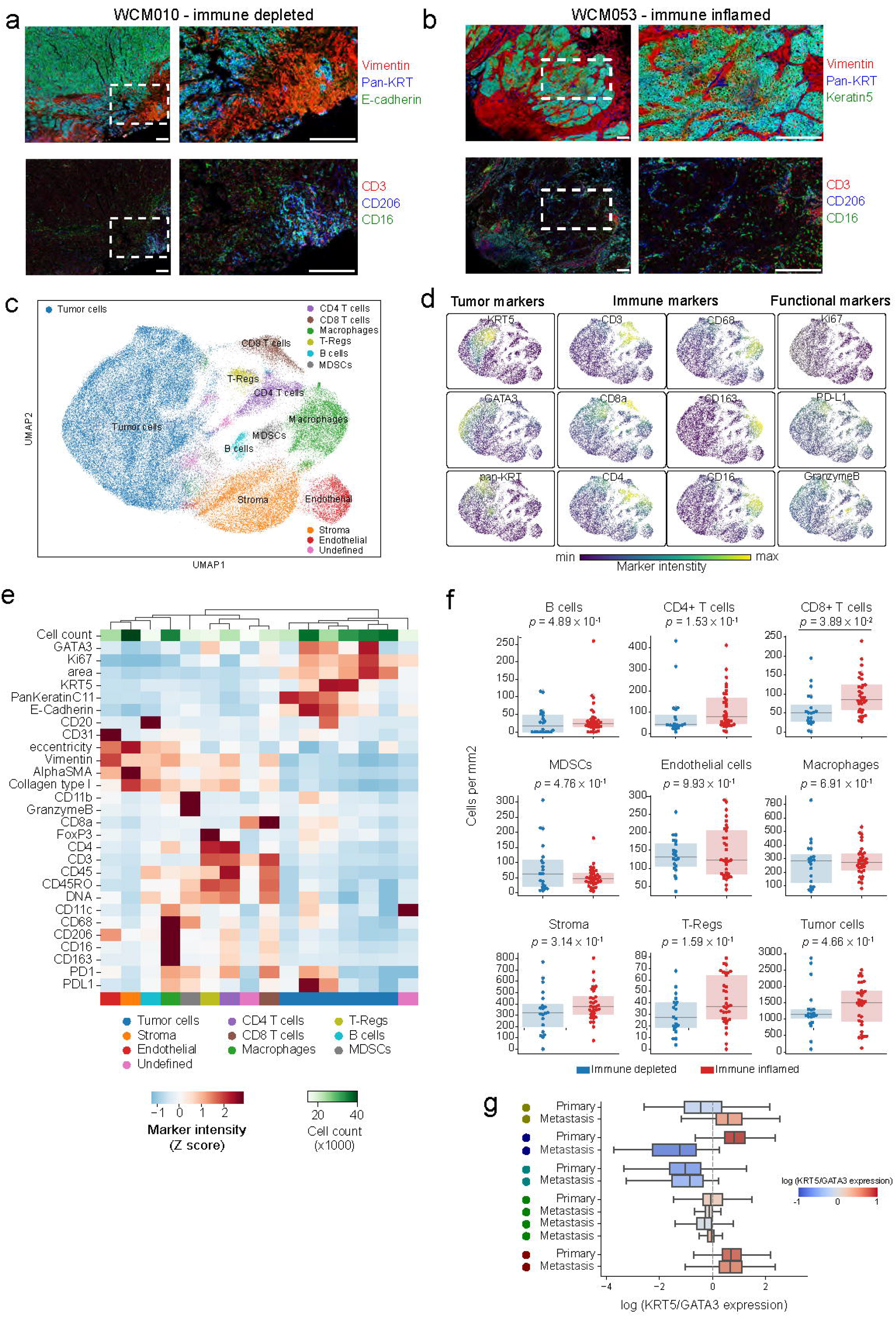
Imaging Mass Cytometry^™^ reveals intra-patient concordance in molecular subtype plasticity between primary and metastatic UTUC tumors at the single cell level. **(a-b)** Representative images of IMC^**™**^ data for a sample of the (a) immune-depleted or (b) immune-inflamed classes. Red: tumor expressed, E-cadherin; Green: T-cell expressed CD3; Blue: DNA. Scale bars represent 200 microns horizontally. **(c)** UMAP representation of single-cells segmented in IMC^**™**^ data colored by the metacluster of origin. **(d)** UMAP representation as in c) but where each single cell is colored by the intensity of various markers in the panel. **(e)** Heatmap of clusters found in the IMC^**™**^ data (columns) and their average intensity in the markers. Clusters were aggregated into meta-clusters dependent on their ontogeny. **(f)** Absolute abundance (normalized to area imaged) of tumor and immune cells dependent on the predicted T-cell depleted or T-cell inflamed phenotypes from RNA-seq. **(g)** Log ratio of KRT5 to GATA3 expression for each single cell aggregated in boxplots for each sample in the IMC dataset. The vertical dotted line represents the level where the KRT5 expression is equal to the GATA3 expression.

## Discussion

Here, we report the genomic, transcriptomic and immunophenotypic landscape and the differences between primary and metastatic UTUC. Furthermore, for the first time, we used multiplexed tissue imaging using Imaging Mass Cytometry^™^, to describe the intra- and inter-tumoral phenotypic heterogeneity of cancer cells and their microenvironment at the single-cell level. Although genomic data from paired primary and metastatic UTUC tumors in small cohorts has been reported^6,20^, neither transcriptomic data nor single-cell imaging analysis have been described.

WES analysis from paired primary and metastatic UTUC revealed that metastatic tumors harbored oncogenic genomic alterations which were not identified in the matched primary tumors. Two possibilities can be considered to account for this heterogeneity: (a) Clones harboring these additional alterations are resistant to chemotherapy. (b) Metastatic lineages result from early branched evolution in the primary tumors. These possibilities were discussed in previous reports which suggested that systemic spread is a very early event in cancer history and that chemotherapy selects clones harboring drug-resistant mutations in breast, colorectal, lung, and bladder cancers^13,21^. In our cohort, intra-patient heterogeneity of likely oncogenic alterations was observed also in the chemotherapy-naïve cases (WCM052 and WCM068). However, the percentages of shared mutations between primary and matched metastases were higher than those who had a history of chemotherapy before sampling. The observation suggests that chemotherapy potentially plays a role in increasing genomic heterogeneity by inducing mutations^13-15^. Regarding UTUC, previous genomic studies in small cohorts showed discordance of mutations between paired primary and metastatic UTUC^6,20^. Furthermore, we show that intra-patient genomic diversity in clinically targetable alterations is also common in UTUC. This finding is of clinical importance because evaluating only a primary or metastatic sites could potentially miss therapeutic targets in patients with metastatic UTUC. However, we acknowledge a limitation that the collected samples may not fully represent the tumors. There is a possibility that some of the alterations detected only in metastases were present in unsampled regions of their paired primary tumors. Liquid biopsy, which can analyze circulating cell-free DNA, could complement tissue biopsy by capturing tumor heterogeneity that single-lesion tumor biopsies may miss^22^.

We observed intra-patient discrepancy of molecular subtypes within 2 patients (WCM01031 and WCM031) at the transcriptomic level using a recently published molecular classification system^17^. A recent study provided important findings which may explain the molecular basis of the molecular subtype shift in urothelial carcinomas. Using a single-cell transcriptome and *in vivo* bladder cancer model, Sfakianos *et al*. revealed lineage plasticity in primary and metastatic urothelial cancers of the bladder^23^. The molecular subtype changes observed in our cohort may reflect the consequence of lineage plasticity during metastatic progression. By contrast, for cases WCM068 and WCM085, molecular subtypes were concordant between matched primary and metastatic tumors. However, we note that intra-tumoral heterogeneity of molecular subtypes may be underestimated due to a sampling bias from a single tumor biopsy, as intra-tumoral and intra-individual molecular heterogeneity has been shown in urothelial carcinomas by previous genomic and transcriptomic studies^24-26^. The intra-individual heterogeneity of molecular subtypes is of potential clinical relevance because molecular classification is expected to predict treatment response and overall survival^27,28^. A single tumor sampling may not reflect the biological characteristic of the disease in the setting of metastatic UTUC.

Contrary to the heterogeneity at genomic levels, the immune phenotype based on the expression of immune-related genes was maintained through the metastatic process. Characterizing the immune microenvironment potentially provides clinical benefit because the extent of tumor-infiltrating T-lymphocytes is positively associated with the efficacy of immune checkpoint inhibitors^29^. Our finding on the consistency of immune phenotype within each patient suggests that evaluating a single tissue site provides a reasonable assessment of the immune microenvironment and thus may be useful for predicting the efficacy of immune checkpoint inhibitors.

Multiplexed imaging using IMC^™^ allowed us to comprehensively profile tumor phenotypes and the microenvironment in primary and metastatic UTUC, revealing the extensive heterogeneity between patients but conserved between primary and metastatic samples of the same individual in all but one patient. While this analysis includes a limited set of protein markers compared with RNA-seq, it allowed us to characterize the spatial heterogeneity of cancer, stromal and immune cells UTUC evolution at the single-cell level. In addition, this enabled us to identify mixed tumor phenotypes within the same sample for several patients. Furthermore, this technique allowed us to validate transcriptome-based predictions of immune inflamed or depleted microenvironment. This spatially-resolved imaging technique generated additional insights not only at the level of quantifying the density of cellular components of the immune compartment such as CD8+ T-cells but also pinpointing their distribution, including the degree of infiltration into the tumor mass compared to their presence in the periphery.

Altogether, our genomic, transcriptomic, and phenotyping analysis of primary and metastatic UTUC brings to light the range of mutational and phenotypic diversity between and within individual patients, revealing a broadly conserved phenotypic tumor landscape against the backdrop of genetic evolution seen during metastasis. This integrated characterization of UTUC informs the targeted and immune therapeutic strategies that maximize efficacy in patients with metastatic UTUC.

## Methods

### Patient enrollment and tissue acquisition

Patients with primary and metastatic UTUC were prospectively enrolled in an institutional review board (IRB)–approved Research for Precision Medicine Study (IRB No. 1305013903) with written informed consent. Tumor tissue from biopsies and nephroureterectomy specimens was collected from 44 patients diagnosed with high-grade urothelial carcinoma. WES data from 7 primary tumor samples presented in this manuscript have already been published^7^. Tumor DNA for WES was obtained from fresh frozen or formalin-fixed, paraffin-embedded tissue. Samples were selected based on pathologic diagnosis according to standard guidelines for UTUC^2,30^. Small cell carcinoma was excluded from our cohort. Pathological review by study pathologists (B.D.R., F.K., J.M.M.) confirmed the diagnosis and determined tumor content.

### DNA extraction and WES

WES was performed on each patient’s tumor/matched germline DNA pair using previously described protocols^31,32^. After macrodissection of target lesions, tumor DNA was extracted from formalin-fixed, paraffin-embedded (FFPE) or cored OCT-cryopreserved tumors using the Promega Maxwell 16 MDx (Promega, Madison, WI, USA). Germline DNA was extracted from blood, buccal mucosa and normal lung and lymph node tissue using the same method. A minimum of 200 ng of DNA was used for WES. DNA quality was determined by TapeStation Instrument (Agilent Technologies, Santa Clara, CA) and was confirmed by real-time PCR before sequencing. Sequencing was performed using Illumina HiSeq 2500 (2 × 100 bp). A total of 21,522 genes were analyzed with an average coverage of 85× using Agilent HaloPlex Exome (Agilent Technologies, Santa Clara, CA).

### WES data processing pipeline

All the sample data were processed through the computational analysis pipeline of the Institute for Precision Medicine at Weill Cornell Medicine and NewYork-Presbyterian (IPM-Exome-pipeline)^31^. Raw reads quality was assessed with FASTQC. Pipeline output includes segment DNA copy number data, somatic copy-number aberrations (CNAs), and putative somatic single-nucleotide variants (SNVs)^32^. Somatic variants were filtered using the following criteria: (a) read depth for both tumor and matched normal samples was ≥30 reads, (b) the variant allele frequency (VAF) in tumor samples was ≥10% and >5 reads harboring the mutated allele, (c) the VAF of matched normal was ≤1%, or there was just one read with mutated allele. Pathogenicity and actionability for each mutation and CNA were determined by the OncoKB database^16^. Tumor mutation burden (TMB) was calculated as the number of mutations divided by the number of bases in the coverage space per million. TMB status (high vs. low) was determined using a urothelial cancer-specific threshold which our group recently reported^10^.

### Computational detection of MSI

MSI was detected by the MSI sensor. MSI sensor is a software tool that quantifies MSI in paired tumor–normal genome sequencing data and reports the somatic status of corresponding microsatellite sites in the human genome^11^. MSIsensor score was calculated by dividing the number of microsatellite unstable by the total number of microsatellite stable (MS) sites detected. The cut-off for defining MSI-high (MSI-H) versus MS stable (MSS) samples was 3.5 (MSI-H > 3.5, MSS < 3.5)^11^.

### RNA extraction, RNA sequencing, and data analysis

RNA-seq and data processing was performed according to the protocol described in previous papers^16,33^. Briefly, RNA was extracted from frozen material for RNA-seq using Promega Maxwell 16 MDx instrument, (Maxwell 16 LEV simplyRNA Tissue Kit (cat. # AS1280)). Specimens were prepared for RNA sequencing using TruSeq RNA Library Preparation Kit v2 or riboZero as previously described^33^. RNA integrity was verified using the Agilent Bioanalyzer 2100 (Agilent Technologies). cDNA was synthesized from total RNA using Superscript III (Invitrogen). Sequencing was then performed on GAII, HiSeq 2000, or HiSeq 2500 as paired-ends^16,33^. All reads were independently aligned with STAR_2.4.0f1^34^ for sequence alignment against the human genome sequence build hg19, downloaded via the UCSC genome browser (http://hgdownload.soe.ucsc.edu/goldenPath/hg19/bigZips/), and SAMTOOLS v0.1.19^35^ for sorting and indexing reads. Cufflinks (2.0.2) ^36^ was used to estimate the expression values (FPKMS), and GENCODE v23^37^ GTF file for annotation. Rstudio with R (v3.6.1) was used for the statistical analysis and the generation of figures.

### Molecular subtyping

The log-transformed expression data was used to infer molecular subtypes using a recently published classification system as implemented in the consensusMIBC R package^17^. Default parameters were set except for minCor representing a minimal threshold for best Pearson’s correlation (minCor = 0.15). The consensus classification implements a nearest centroid method and Pearson’s correlation and classifies samples into 6 molecular classes; Luminal Papillary (LumP), Luminal Non Specified (LumNS), Luminal Unstable (LumU), Stroma-rich, basal/Squamous (Ba/Sq), Neuroendocrine-like (NE-like).

### T cell inflammation classification analysis

The previously published T cell inflammation gene signature was used to classify tumors into T-cell inflamed or T cell depleted^7^. The expression data, quantified as FPKMs, was obtained for the EIPM UTUC patients, of which RNA-seq was available (n=17). The FPKMs for the primary tumors were obtained from the GDC/TCGA bladder cohort (TCGA BLCA, n=414). The genes from the signature were selected for expression-based supervised clustering. The FPKMs were log-transformed and median centered, and partitioning around medoids (PAM) algorithm was applied to cluster the transformed expression data. This led to the identification of two broad clusters: one with higher expression of the signature genes was labeled as T cell inflamed while that with low expression of signatures genes was labeled as T cell-depleted.

### Imaging Mass Cytometry^™^

Antibodies were conjugated in BSA and Azide free format using the MaxPar X8 multimetal labeling kit (Fluidigm) as per the manufacturer’s protocol. Freshly cut 4-micron thick FFPE tissue sections were stored at 4oC for a day before staining. Slides were first incubated for 1 hour at 60oC on a slide warmer followed by dewaxing in fresh CitriSolv (Decon Labs) twice for 10 minutes, rehydrated in descending series of 100%, 95%, 80%, and 75% ethanol for 5 minutes each. After 5 minutes of MilliQ water wash, slides were treated with antigen retrieval solution (Tris-EDTA pH 9.2) for 30 minutes at 96oC, cooled to room temperature (RT), washed twice in TBS, and blocked for 1.5 hours in SuperBlock Solution (ThermoFischer). Overnight incubation occurred at 4oC with the prepared antibody cocktail containing the metal-labeled antibodies. The next day, slides were washed twice in 0.2% Triton X-100 in PBS, and twice in TBS. DNA staining was performed using Intercalator-Iridium in PBS solution for 30 minutes in a humid chamber at room temperature, followed by a washing step in MilliQ water and air drying. The Hyperion instrument was calibrated using a tuning slide to optimize the sensitivity of the detection range. Hematoxylin and Eosin (H&E) stained slides were used to guide the selection of regions of interest in order to obtain representative regions. All ablations were performed with a laser frequency of 200 Hz. Tuning was performed intermittently to ensure the signal detection integrity was within the detectable range.

### Analysis of Imaging Mass Cytometry^™^ data

Imaging Mass Cytometry^™^ data were preprocessed as previously described^9^ with some modifications. Briefly, image data was extracted from MCD files acquired with the Hyperion instrument. Hot pixels were removed using a fixed threshold. The image was amplified two times, Gaussian smoothing applied and, from each image, a square 500-pixel crop was saved as an HDF5 file for image segmentation. Segmentation of cells in the image was performed with *ilastik* (version 1.3.3) ^38^ by manually labeling pixels as belonging to one of three classes: nuclei – the area marked by a signal in the DNA and Histone H3 channels; cytoplasm – the area immediately surrounding the nuclei and overlapping with signal in cytoplasmic channels; and background – pixels with low signal across all channels. Ilastik learns from these sparse labels by training a Random Forest classifier using features present in the images. Features used were the Gaussian Smoothing with kernel widths of 1 and 10 pixels, Hessian of Gaussian Eigenvalues with kernel 3.5 and 10 pixels, and Structure of Tensor Eigenvalues with kernel of 10 pixels. The outputs of prediction are class probabilities for each pixel which were used to segment the using DeepCell version 0.8.2)^39^ with the *MultiplexSegmentation* pre-trained model.

To identify cell types in an unsupervised fashion, we first quantified the intensity of all samples in each segmented cell, not overlapping image borders. Channels with “functional” markers Ki67, PD-1, PD-L1, and Granzyme B were not used for downstream cell type identification but only for visualization. In addition, for each cell, we computed morphological features such as the cell area, perimeter, the length of its major axis, eccentricity, and solidity using the skimage.measure.regionprops_table function (version 0.18.1)^39^. Cells with solidity value of 1 (perfectly round cells) were excluded from analysis. Using Scanpy (version 1.7.1)^40^, we log-transformed the quantification matrix, and Z-scored values per image, capping the signal at - 3 and 3, followed by global feature centering and scaling. The batch was removed with Combat (Scanpy implementation) and features scaled again. Principal Component Analysis was performed, and we computed a neighbor graph on the PCA latent space using the batch balanced k-nearest neighbors (bbknn) (version 1.4.0)^41^. We computed a Uniform Manifold Approximation and Projection (UMAP)^42^ embedding (umap package, version 0.4.6) with a gamma parameter of 25, and clustered the cells with the Leiden algorithm^43^ with resolution 0.5 (leidenalg package, version 0.8.3).

### Statistical analyses

For statistical tests, a two-sided Mann–Whitney test was used to check for significant differences between two distributions. The two-sided Fisher’s exact test was applied to determine whether the deviations between the observed and the expected counts were significant. We used a *P* value threshold of 0.05.

## Acknowledgments

This work was supported by the Cornell Center for Immunology Core Facilities Seed Grant (PI: J.M.M., Co-PI: K.O., and Co-I: O.E., B.M.F.), and by the Caryl and Englander Institute for Precision Medicine. A.F.R. is supported by an NCI T32CA203702 grant. B.M.F. was supported by the Department of Defense CDMRP grant CA160212 and Gellert Family-John P. Leonard, MD Research Scholarship in Hematology and Medical Oncology. For technical support, the authors thank Bing He, Ruben Diaz, Leticia Dizon from the Center for Translational Pathology of the Department of Pathology and Laboratory Medicine at Weill Cornell Medicine.

## Author contributions

Initiation and design of the study: K.O., B.M.F. and J.M.M. Subject enrollment, sample, pathology review and clinical data collection: K.O., A.V., S.B., B.D.R., F.K., B.M.F. and J.M.M. Lab data collection and analysis: K.O., W.L., K.L. and B.M.F. Imaging mass cytometry: H.R. and statistical and bioinformatic analyses: K.O., B.B., K.W.E., E.V., R.B., A.F.R., A.S. and O.E. Supervision of research: B.M.F. and J.M.M. Writing of the first draft of the manuscript: K.O., B.M.F. and J.M.M. All authors contributed to the writing and editing of the revised manuscript and approved the manuscript.

## Competing interests

B.M.F. has received research support for Weill Cornell from Eli Lilly and served on advisory boards for Immunomedics, Seattle Genetics, QED Therapeutics, Merck & Co. Consulted for QED Therapeutics, received patent royalties from Immunomedics and Gilead Sciences, and received honoraria from Urotoday.

## Notes

### Competing Interest Statement

The authors have declared no competing interest.

