## Supplementary material for "The evolution of genomic, transcriptomic, and single-cell protein markers of metastatic upper tract urothelial carcinoma": Supplmentary figures and legend

### Supplementary Figure 1

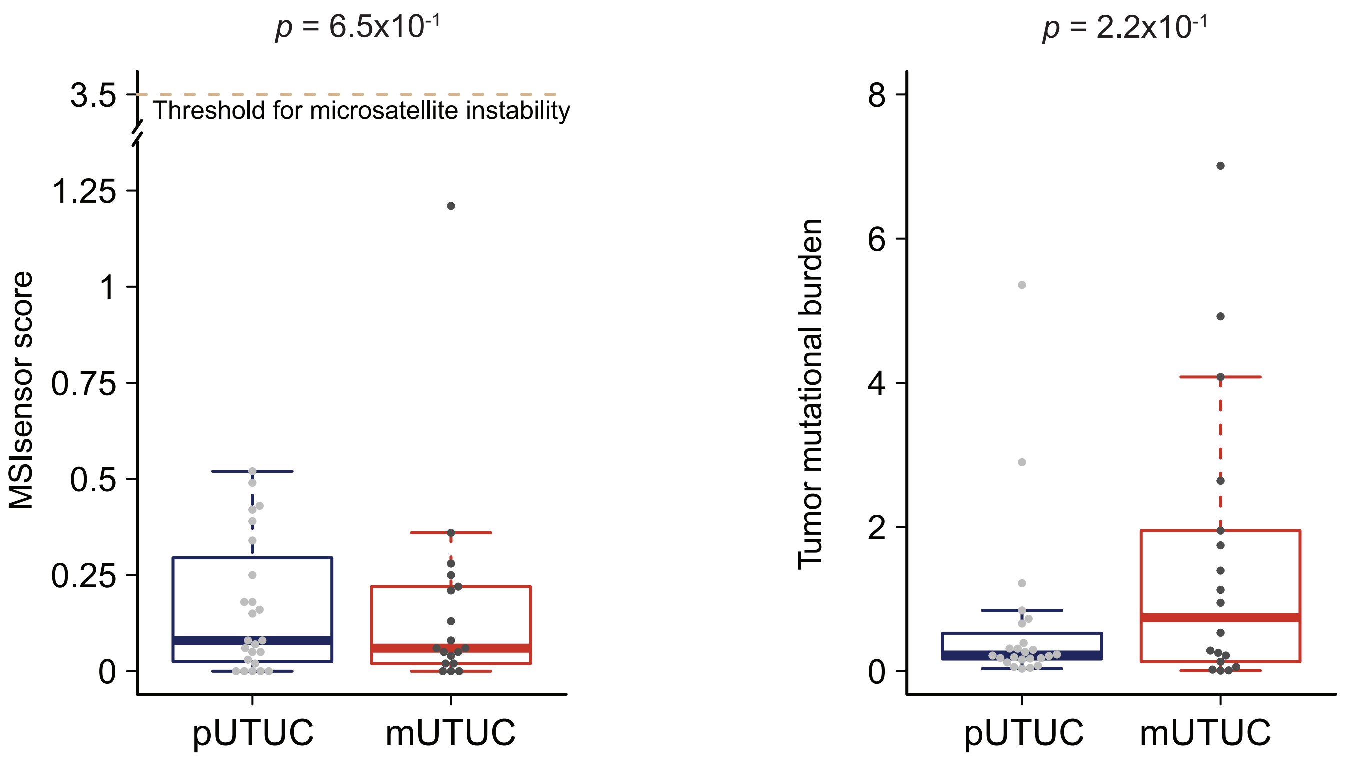

### Supplementary Figure 2

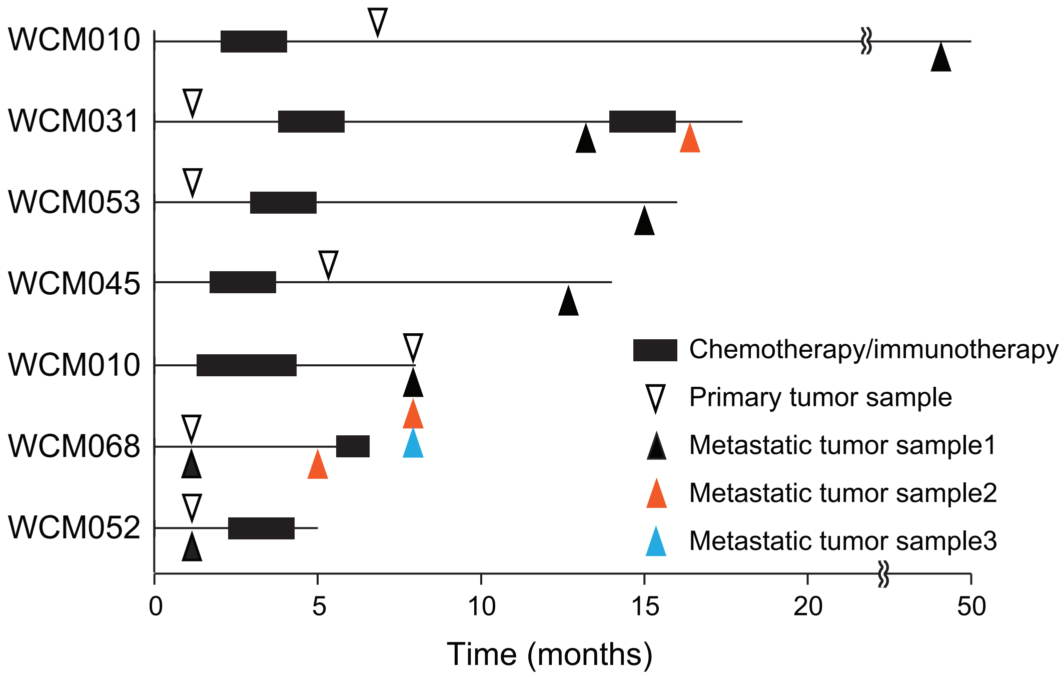

### Supplementary Figure 3

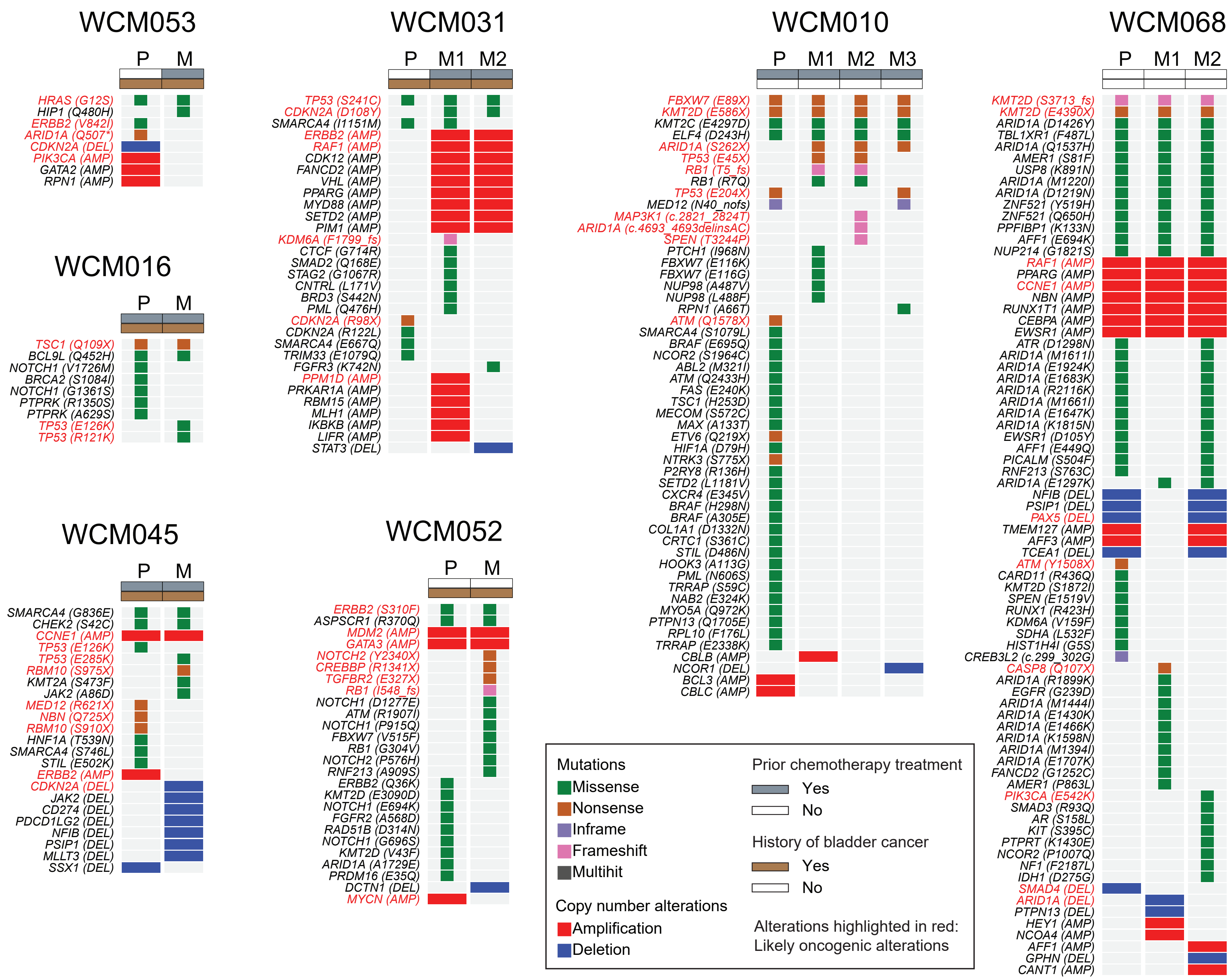

### Supplementary Figure 4

WCM031

Primary Metastasis

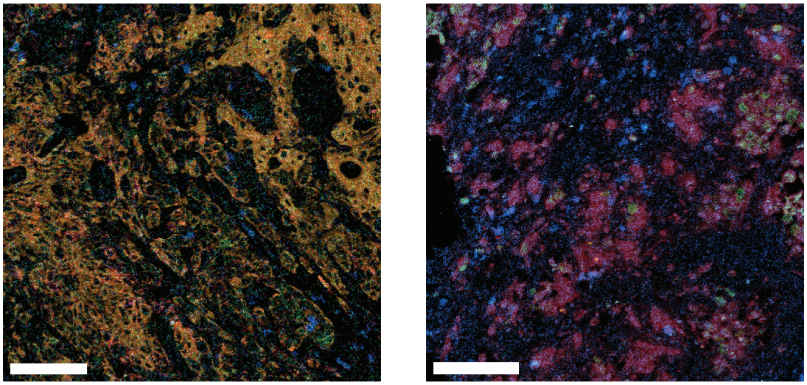

CD3 KRT5 GATA3 pan-KRT

| | Primary | Metastasis |
|---|---|---|
| Ba/Sq vs Lum (log ratio) | 0.83 | -1.25 |
| CD8 T cells (per mm2) | 78.6 | 134.5 |
| Phenotype predicted | Ba/Sq, Depleted | Lum, Inflammed |

Supplementary Figure 1. Comparison of microsatellite instability (MSI) and total mutational burden (TMB) between primary and metastatic UTUC. No significant differences are shwn in MSIsensor scores and TMB between primary and metastatic UTUC.

Supplementary Figure 2. Timeline and clinical course (vertical lines) of patients with paired primary and metastatic UTUC.

Supplementary Figure 3. Somatic mutations, amplification, and deletion of cancer genes in primary (P) and matched metastatic (M) UTUC. Discordance in likely oncogenic alterations between primary tumor and paired metastases is shown.

Supplementary Figure 4. Regions of interest for samples of patient WCM031 illustrating differential tumor phenotype and immune infiltration between primary and metastasis. Primary tumor cells show cytoplasmic staining of the basal/squamous (Ba/Sq) marker KRT5. Metastatic tumor cells demonstrate nuclear staining of the luminal (Lum) marker GATA3. CD3 staining highlights low T cell infiltration (depleted) in the primary tumor and high T cell infiltration (inflamed) in the metastatic tumor. Scale bar, 100 µm.
